# Single-cell-resolved dynamics of chromatin architecture delineate cell and regulatory states in wildtype and *cloche/npas4l* mutant zebrafish embryos

**DOI:** 10.1101/2020.06.26.173377

**Authors:** Alison C. McGarvey, Wolfgang Kopp, Dubravka Vučićević, Rieke Kempfer, Kenny Mattonet, Antje Hirsekorn, Ilija Bilić, Alexandra Trinks, Anne Margarete Merks, Daniela Panáková, Ana Pombo, Altuna Akalin, Jan Philipp Junker, Didier Y.R. Stainier, David Garfield, Uwe Ohler, Scott Allen Lacadie

## Abstract

DNA accessibility of cis regulatory elements (CREs) dictates transcriptional activity and drives cell differentiation during development. While many of the genes that regulate embryonic development have been described, the underlying CRE dynamics controlling their expression remain largely unknown. To address this, we applied single-cell combinatorial indexing ATAC-seq (sci-ATAC-seq) to whole 24 hours post fertilization (hpf) stage zebrafish embryos and developed a new computational tool, ScregSeg, that selects informative genome segments and classifies complex accessibility dynamics. We integrated the ScregSeg output with bulk measurements for histone post-translational modifications and 3D genome organization, expanding knowledge of regulatory principles between chromatin modalities. Sci-ATAC-seq profiling of *npas4l*/*cloche* mutant embryos revealed novel cellular roles for this hemato-vascular transcriptional master regulator and suggests an intricate mechanism regulating its expression. Our work constitutes a valuable resource for future studies in developmental, molecular, and computational biology.

## Introduction

The coordination of cis-regulatory elements (CREs) is essential to the tight regulation of gene expression programs that direct cell fate changes in embryonic development. Types of CREs include promoters, enhancers, insulators and silencers, whose sequence and dynamic physical properties determine their function. The fundamental unit of a CRE is a nucleosome-depleted region (NDR) which acts as a binding platform for transcriptional regulators and can be highly dynamic across cell types due to the combined action of pioneering factors and nucleosome remodelers. Mammalian NDRs often harbor divergently-oriented core promoter sequence elements and transcription initiation, and are flanked by well-positioned nucleosomes whose histone post-translational modifications (PTMs) reflect the activation state and/or class of CRE^1,2^. Taken together, this complex architecture has been described as the regulatory interface between the genome and its functional output^3^.

The development of single-cell high-throughput molecular assays^4–6^ has revolutionized systems genomics, allowing the extensive profiling of cell type diversity making up almost any tissue or organism with little to no prior information. The Assay for Transposase-Accessible Chromatin using sequencing (ATAC-seq)^7^ can quantify the extent of CRE nucleosome depletion at a genome-wide scale. Its further development has enabled the measure of chromatin accessibility in tens of thousands of single cells from diverse biological contexts such as: cell lines^8,9^, mouse primary tissues^10–13^, *Drosophila* embryos^14^, human primary tissues^15,16^ and human organoids^17^. These datasets have generated comprehensive resources of putative distal regulatory elements, transcriptional regulators, cell type specificity of inherited disease associated traits^10,15^, putative higher-order interactions between regulatory elements^9^, as well as epigenomic contribution to lineage priming^16^. However, sc-ATAC-seq data presents several analysis challenges distinct from sc-RNA-seq measurements, therefore computational method development for this data type is an important and ongoing issue.

Zebrafish has a long history as a model system for embryology, and forward genetic screens have identified many genes with key roles during vertebrate development. More recently, zebrafish have been increasingly used for cutting-edge genomic profiling^18–23^, however, cis-regulatory dynamics have yet to be characterized at single-cell resolution. Furthermore, key resources available for mouse or human genomics studies, such as genome classifications based on histone PTM ChIP-seq signals, high-depth genome-wide probing of 3D chromatin spatial organization, and databases of regulatory elements are limited for the zebrafish community.

Here we characterized the genome-wide chromatin architecture of the whole 24 hpf stage zebrafish embryo, at bulk and single cell resolution, to generate a resource of cell type-specific candidate CREs. We applied sci-ATAC-seq^14^ to whole embryos, generating accessibility profiles for ∼23,000 single nuclei. We developed a tool, named ScregSeg, which utilizes Hidden Markov Models (HMMs) to address key challenges in analyzing single-cell accessibility profiles: 1) the selection of informative regions for the downstream analysis, and 2) the characterization of complex cell-specific CRE dynamics. We show that diverse cell types present in the 24 hpf embryo can be identified by their accessibility profiles and have classified complex patterns of CRE dynamics that reflect the combinatorial nature of transcriptional regulation. Sequence analysis of these cis-regulators allowed us to infer putative transcription factors that bind chromatin in a cell type-specific manner. Using bulk ChIP-seq data for histone PTMs known to occur at CREs, we provide the additional resource of a genome-wide classification for promoter- and enhancer-like chromatin states at the 24 hpf stage. Integrating these classifications with sci-ATAC-seq and bulk *in situ* Hi-C, we show clear relationships between promoter-like states, constitutive accessibility, and 3D insulation, as well as between co-accessibility and 3D interactions, thereby expanding insight into regulatory principles active during zebrafish development. Lastly, we apply sci-ATAC-seq to embryos harboring a mutation in the *cloche* gene, *npas4l*, which lack blood and endothelial cells^24,25^, and observe hitherto undescribed changes in muscle, epidermal, and caudal precursor cell numbers^24,25^. Furthermore, we detect candidate cell type-specific *npas4l* CREs, suggesting an intricate network controlling this hemato-vascular transcriptional master regulator.

## Results

### Single-nucleus accessibility profiles separate whole embryos into cell types

We set out to determine a comprehensive genomic regulatory map of the zebrafish, *Danio rerio*, at the 24 hpf stage with the bipartite goal of 1) uncovering regulatory architecture and principles, and 2) establishing a resource of genome-wide regulatory classifications for future studies. At 24 hpf, zebrafish embryos have established the classic bilateral vertebrate body plan and are in a key transitional point of cell type specification and organogenesis, arguably the most morphologically comparable across diverse vertebrate embryos^26,27^. DNA accessibility within chromatin is highly dynamic between cell types^28^, mostly reflecting regulatory changes. We, therefore, employed single-nucleus combinatorial indexing^14^ to determine genome-wide accessibility profiles for many cell types in parallel from whole zebrafish embryos. Nuclei were isolated from staged embryos, subjected to two rounds of barcoding via tagmentation and PCR with random mixing in between, and the resulting DNA fragments sequenced to high-depth (Figure 1a). Species-mixing with nuclei from the sea urchin demonstrated single-cell resolution with an acceptable barcode-collision rate of ∼14-15%^29^, and a distinct distribution of barcodes with > 1000 unique reads was considered to represent intact nuclei (Figure S1a,b). In all, we sequenced approximately 23,000 nuclei with an average depth per nucleus of more than 10,000 from three independent experiments: two wildtype experiments and one from a mixture of wildtype and embryos harboring a mutation in the critical developmental transcription factor, *npas4l*, otherwise known as *cloche* (see below for detailed characterization).

**Figure 1.**
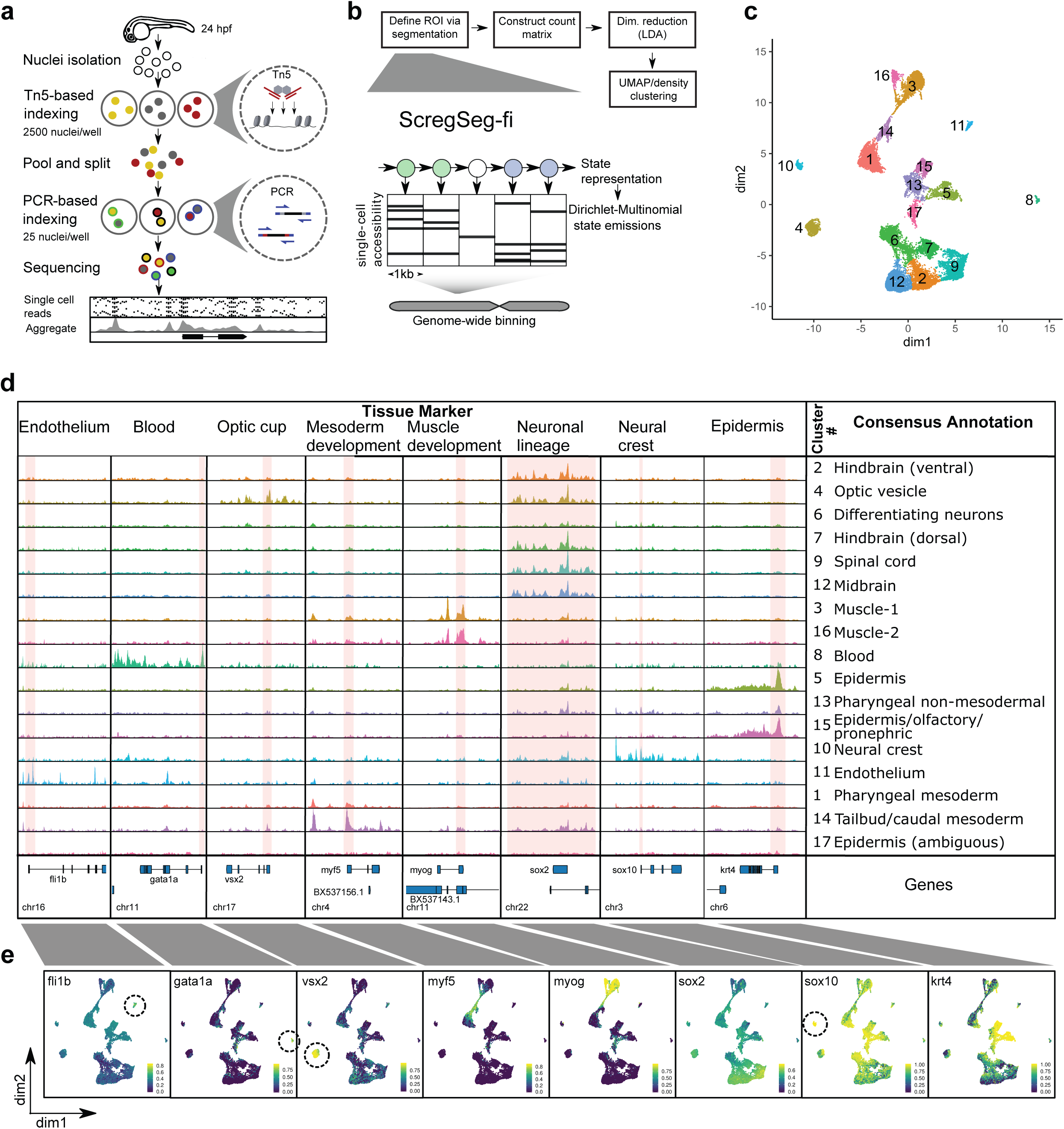
Generating cell type specific accessibility profiles from 24 hpf zebrafish embryos. a. Schematic of sci-ATAC-seq method. Nuclei are extracted from flash frozen whole embryos staged at 24 hpf. Nuclei are sorted into 96-well plates, 2500 per well, and barcoded during tagmentation. Tagmented nuclei are pooled then split into 96-well plates, 25 per well, and a second set of barcodes introduced by PCR. The resulting DNA fragments are pooled and sequenced, with unique barcode combinations representing single cells. b. Schematic representation of ScregSeg for genomic feature identification (ScregSeg-fi) and clustering of single cell accessibility profiles. The genome is divided into 1 kb bins and a HMM-based segmentation is used to assign a state to each bin based on the accessibility distribution over cells. Informative states are used to construct a cell by genomic region count matrix, which undergoes dimensionality reduction using LDA implemented in cisTopic, batch correction, UMAP analysis and density clustering. c. UMAP representation of the cell-Topic matrix from cisTopic on ∼23000 cells from three experiments. Colors represent the 17 clusters determined by density clustering. d. Summary pseudo-bulk chromatin accessibility profiles from aggregated cells for each density cluster at marker genes of major tissues and cell types of 24 hpf zebrafish embryos. Consensus annotations derived from enrichment of genes that map to differentially accessible segments per cluster, with ZFIN anatomical database terms, and published cell-type markers ^23^. Highlighted regions are displayed in E. e. Per cell distribution of accessibility at regions covering the promoters of marker genes (highlighted in D), represented in UMAP space. Color represents the rank-based AUCell enrichment score for a given region^30,95^.

Whereas single-cell RNA expression measurements are typically quantified based on known gene annotations, the initial definition of features to be quantified from single-nucleus accessibility maps poses a significant challenge. Two current approaches to this problem include peak calling on the bulk data or genome-wide binning. Peak calling has the disadvantage of missing highly cell-specific regions associated with small cell populations, while genome-wide binning leads to a large number of uninformative features which may overwhelm downstream computations. Here, we employed a novel strategy whereby single-nuclei signal tracks in genome-wide 1 kb resolution are subjected to a Hidden Markov Model (HMM)-based segmentation with Dirichlet-Multinomial state emissions. The model describes the relative cross-cell accessibility profiles using different states, which capture cell type-specific regulatory activity while also accounting for correlations between neighboring genomic regions. The states can be broadly divided into abundant and rare, the former reflecting background signal, the latter cell type-specific accessibility profiles. Rare states tend to show 1) higher read coverage across the cells, 2) higher ChIP-seq signal for the CRE-associated histone PTM H3K27ac, and 3) enrichment for short fragments associated with NDRs (Figure S1c-g). Therefore, we selected a set of rare states, which together represent diverse accessibility profiles (e.g. regions representing variable accessibility dynamics), to define the informative genomic regions for downstream analysis. We call this approach <u>s</u>ingle-<u>c</u>ell <u>reg</u>ulatory landscape <u>seg</u>mentation for feature identification (ScregSeg-fi).

Having identified informative features (segments) we performed dimensionality reduction with Latent Dirichlet Allocation (LDA) using cisTopic^30^ (Figure 1b, S1h). The resulting low dimensional matrix was subjected to batch correction followed by UMAP transformation. Finally, grouping nuclei according to density in the UMAP space lead to 17 clusters as candidate cell types (Figure 1c, S1i; see Methods).

To annotate the sci-ATAC-seq clusters, we performed differential analysis of the informative segments identified by ScregSeg-fi to identify marker regions unique to each cluster (see Methods). We assigned these regions to the closest annotated gene transcription start site. We tested enrichment of these marker genes within gene sets from the ZFIN anatomical database and published stage-matched single-cell RNA-seq^23,31^ (Tables S1, S2) and derived consensus annotations with variable confidence (Figure 1d). Endothelium, blood, neural crest, and optic vesicle annotations could be confidently assigned to four distinct clusters (Figure 1c; clusters 11, 8, 10, 4, respectively), showing accessibility around known marker genes, *fli1b*^*32*^, *gata1a*^*33,34*^, *sox10*^*35*^ and *vsx2*^*36*^, respectively (Figure 1d-e). Three, more distributed, territories with substructure were also observed: 1) A largely mesodermal territory encompassing: clusters 16 and 3, which show high accessibility around muscle cell marker genes such as *myog*; cluster 14, which has high accessibility around early mesoderm markers such as *myf5*, presumably representing less differentiated tailbud precursors and caudal mesoderm; and cluster 1, which is enriched with pharyngeal mesoderm markers. The division of muscle into clusters 16 and 3 is likely due to differences in cells harboring the *npas4l* mutation, described below in detail (Figure S1i; Figure 5; see Methods). 2) In the center of the map, cluster 5 and cluster 15 have high accessibility around epidermal and peridermal markers such as *krt4*, and additionally in cluster 15, “olfactory placode” and “pronephric duct” annotations; cluster 13 is enriched for “pharyngeal arch” and “skeletal” terms, and given its location in the UMAP space, is likely to represent non-mesodermal contribution to the pharyngeal arches; cluster 17 which, despite showing low but significant enrichment for epidermal terms, displays a relatively flat accessibility profile suggesting it may represent a different cell state such as mitotic or dying cells, although no such GO terms were enriched (data not shown). 3) A largely neuronal territory with broad enrichment around the early neuronal regulator *sox2*^*37*^ that is subdivided into: cluster 9, spinal cord; cluster 7 and cluster 2 representing hindbrain; cluster 6, representing differentiating neurons; and cluster 12, representing midbrain (Figure 1c-e, Table S1). Based on these marker gene associations, the clusters are, from here on, assigned representative names (Figure 1d).

**Figure 2.**
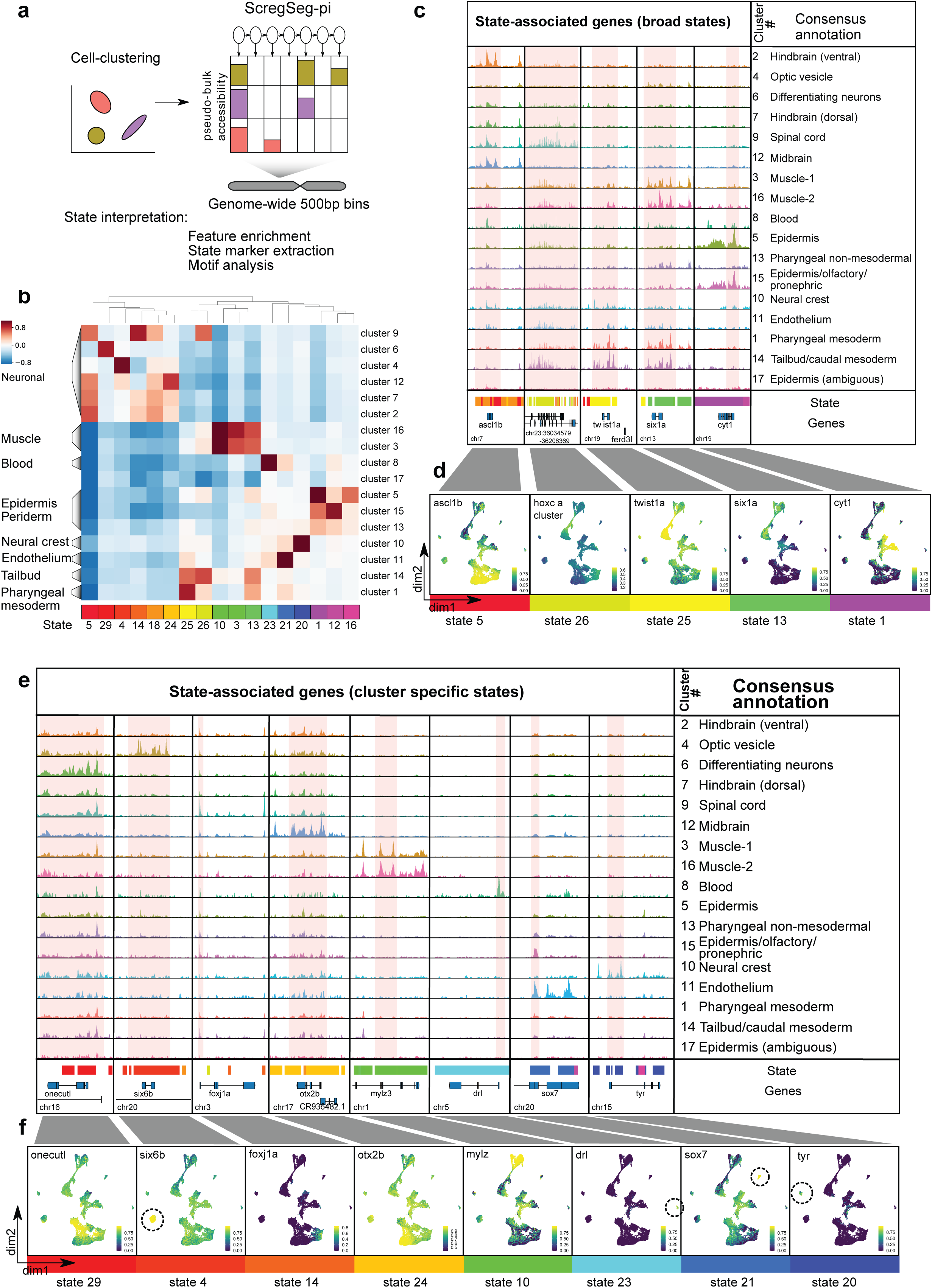
Segmentation of accessibility profiles reveals cell-type specific and shared regulatory regions. a. Schematic representation of ScregSeg for identifying regulatory programs (ScregSeg-pi). The genome is divided into 500 bp bins and a HMM-based segmentation is used to assign a state to each bin based on the accessibility distribution over the density cluster pseudo-bulk accessibility tracks. b. Heat map representing the association between states and clusters based on the log-fold enrichment between the states’ emission probabilities and the genomic background coverage profile (see Methods). Display restricted to states showing the strongest association with clusters (full heatmap in supplementary figure 2A). c. Representative genomic loci specific to broad cluster-associated states 5, 26, 25, 13, or 1. Chromatin accessibility tracks are from aggregated cells from each cluster. Highlighted regions are displayed in D. d. Per cell distribution of accessibility at regions covering the promoters of marker genes (highlighted in C), represented in UMAP space. Color represents the rank-based AUCell enrichment score for a given region ^30,95^. e. Representative genomic loci specific to cluster-specific states 29, 4, 14, 24, 10, 23, 21, or 20. Chromatin accessibility tracks are from aggregated cells from each cluster. Highlighted regions are displayed in F. f. Per cell distribution of accessibility at regions covering the promoters of marker genes (highlighted in E), represented in UMAP space (enrichment score coloring as in D).

**Figure 3.**
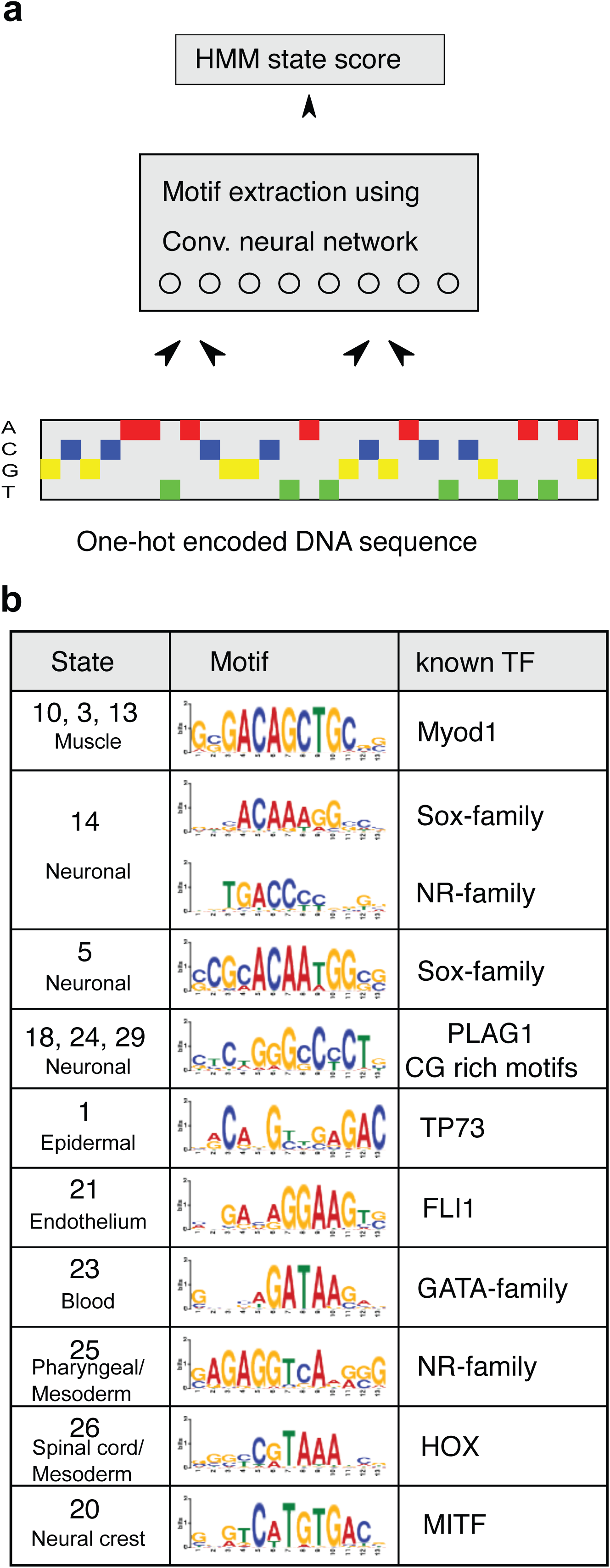
Motif extraction via deep learning. a. A convolutional neural network was employed to extract sequence motifs that are predictive of the expected state-read depth score, a combination of the segmentation model’s state calls and the read depth across cells (see Methods). b. Extracted motifs agree with known motifs of transcription factors implicated in regulating distinct cell-type specific processes.

**Figure 4.**
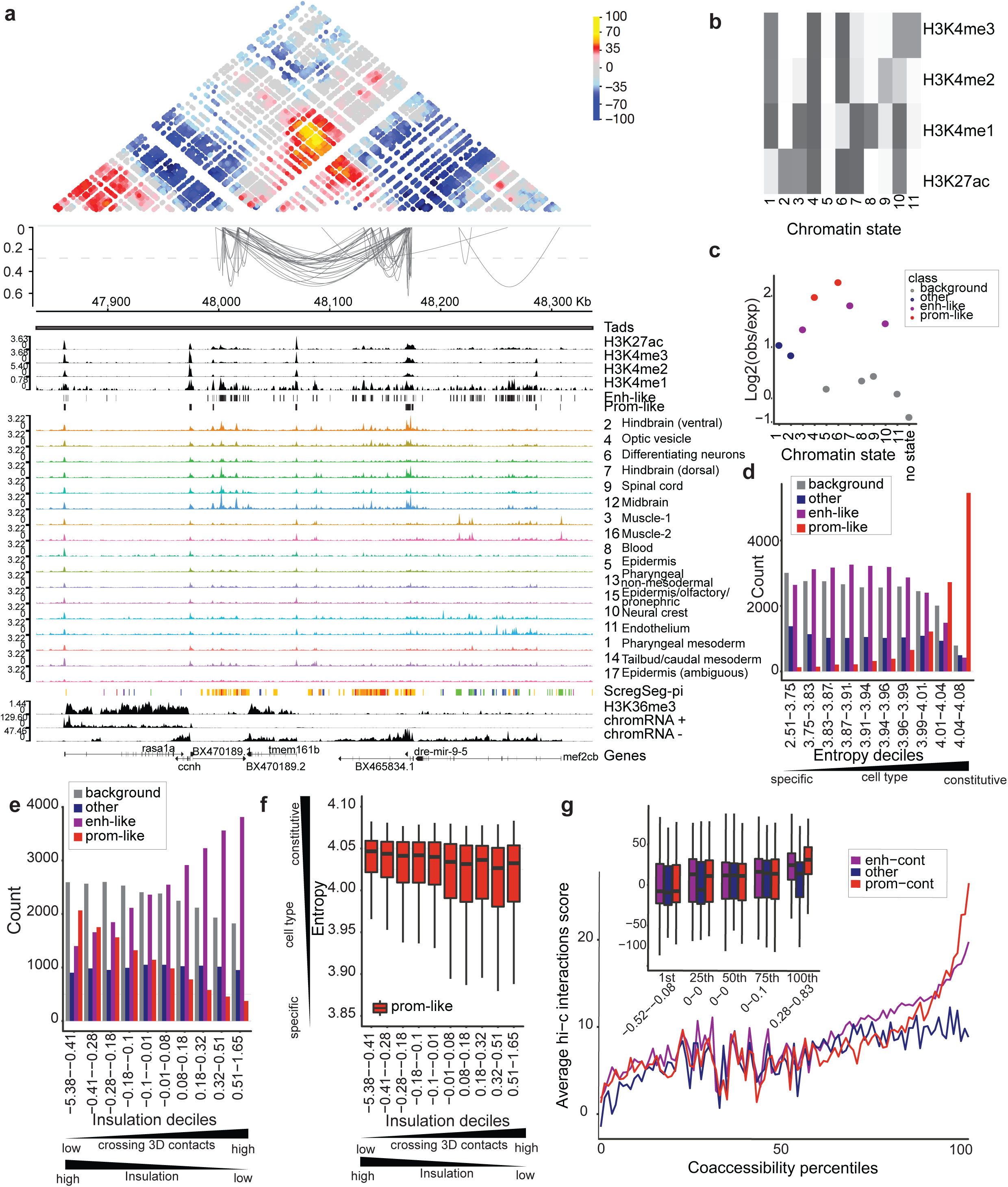
Accessibility dynamics are reflected in histone PTM states and 3D genome organization. a. Browser shot around the dre-mir-9-5 locus showing strong concordance between (from top to bottom) SHAMAN 3D interaction score heatmap, Cicero co-accessibility loops for scores over 0.28 (top 1% score cutoff;dashed line), histone PTM signals and promoter-like/enhancer-like HMM state calls, cluster-collapsed sci-ATAC-seq signals, sci-ATAC-seq segmentation calls, H3K36me3 signal, and nascent chromatin-associated RNA signal. Co-accessibility loops are clearly enriched between strongly interaction regions (orange/yellow in Hi-C heatmap) and these anchor points are clearly marked with enhancer-like and promoter-like PTMs, as captured by the histone PTM states. Co-accessibility is also observed in the sci-ATAC-seq signal tracks and reflected in similar coloring of the sci-ATAC-seq segmentation calls. b. A heatmap representing histone PTM chromatin states learned. Each state is a multivariate Gaussian distribution and is plotted as the mean scaled ChIP-Seq signal for each PTM. c. 1 kb segments from the sci-ATAC-seq foreground are classified for their most representative histone PTM state (see Methods) and plotted is the log2 ratio of class occurrence therein compared to class occurrence in all genomic 1 kb bins. The color scale represents the type of histone PTM state as determined from genome-wide frequency and positional enrichment around annotated-TSS proximal and distal segments (Figure S4B-D). d. Entropy scores (low = cell-specific, high = constitutive) for foreground sci-ATAC-seq segments were split into deciles and within each decile the number of segments for each type of histone PTM state were counted and plotted. e. *In situ* Hi-C insulation scores for foreground sci-ATAC-seq segments were split into deciles and within each decile the number of segments for each type of histone PTM state were counted and plotted. f. *In situ* Hi-C insulation scores for foreground sci-ATAC-seq segments were split into deciles and then split according to their histone PTM type. The entropy score is plotted for the resulting promoter-like histone PTM segments and the other three histone PTM types can be seen in supplemental Figure S4I. g. SHAMAN Hi-C interaction score means (full plot lines) or distributions (inset box plots) are plotted for pairs of sci-ATAC-seq foreground segments that are >25 kb apart and within the same TAD. Segment pairs are split first by Cicero co-accessibility score percentiles and then by having a promoter-like histone PTM state in one or both of the two segments (prom-cont), having no promoter-like histone PTM segments but having one or two enhancer-like PTM segments (enh-cont), or where neither segment is promoter-like or enhancer-like (other). Mean lines for all 100 percentiles are plotted for ease of visualization and box-plot insets for the 1st, 25th, 50th, 75th, or 100th percentiles are shown to give a better sense of the distributions. Counts for each group can be seen in Figure S4J.

**Figure 5.**
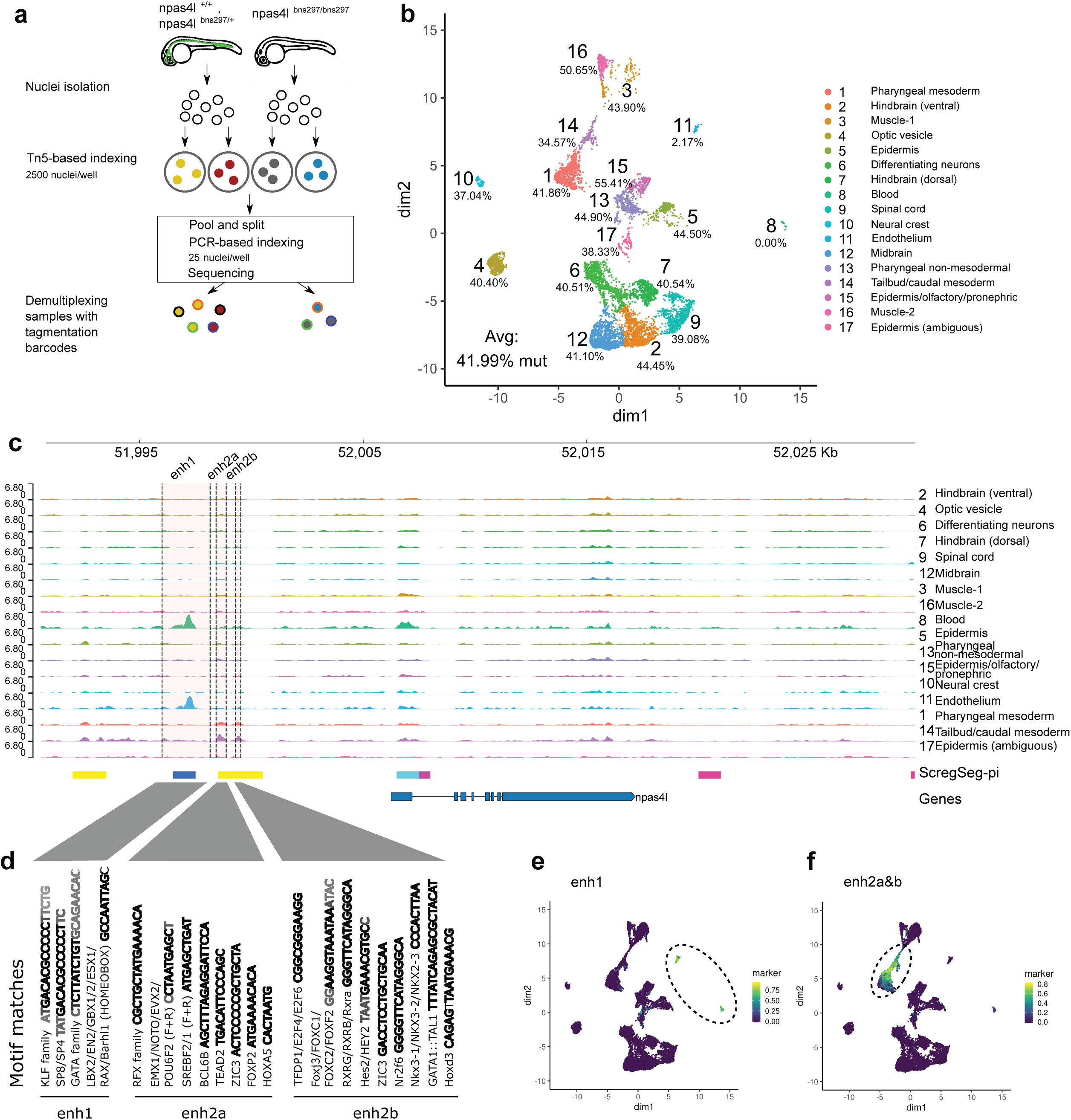
Quantitative analysis of *npas4l* mutant accessibility profiles reveals differences in cell populations and putative novel enhancers. a. Schematic of sci-ATAC-seq data collection from *npas4l* mutant embryos. The Tg(fli1a:nls-GFP)^y7^ background was employed to separate homozygous cloche mutant (*npas4l*^*bns297/bns297*^) embryos from heterozygous and homozygous wild type siblings (*npas4l*^*bns297/+*^, *npas4l*^*+/+*^) based on the loss of a fli1a-GFP+ endothelial cells in the mutant. These two pools of embryos were kept separate for the nuclei isolation and first indexing step (tagmentation). All nuclei were pooled for subsequent steps (as figure 1A) and homozygous mutant vs sibling cells could be distinguished based on their tagmentation barcodes. b. UMAP representation of the cell-Topic matrix from cisTopic on 8976 cells, 3769 homozygous *npas4l* mutant and 5207 siblings. Percentages represent the proportion of mutant cells relative to all mutant and sibling cells per density cluster. c. Summary chromatin accessibility from aggregated cells for each cluster (pseudo-bulks) at the *npas4l* locus. Three cell type-specific peaks of accessibility are highlighted as putative enhancers enh1, enh2a, enh2b around 8-10 kb from the *npas4l* TSS. d. Motif detection in the enh1, enh2a, enh2b sequences with JASPAR motifs^56^. Motif scanning at the specific enhancer was with FIMO^94^ and the 20 highest enriched motif sequences are displayed, collapsed per family. Bold black represents a core sequence match shared across the whole family, and grey represents less frequent variations of the motif sequence. e. Per cell distribution of accessibility at putative *npas4l* enhancer enh1 (highlighted in C), represented in UMAP space. f. Per cell distribution of accessibility at putative *npas4l* enhancer enh2a and enh2b (highlighted in C), represented in UMAP space.

### ScregSeg defines single-cluster- and multi-cluster-specific accessibility dynamics

Cell diversity results from the implementation of regulatory “programs”, which represent unique combinatorial activities of multiple molecular regulators. However, individual components of these programs may be reused in several different contexts. Therefore stringent one-versus-all differential CRE accessibility analysis between single-cell clusters most likely underrepresents the regulatory landscape dynamics due to the subtraction of regions common to multiple cell types. To address this problem, ScregSeg can be utilized by changing the nature of the input matrix (Figure 2a): instead of single-cell signal tracks, we collapsed accessibility counts in genome-wide 500 bp bins for nuclei within the same cell type cluster (Figure 1c), yielding 17 pseudo-bulk signal tracks. The model segments the genome into “states” that describe distinct dynamics across the cell type clusters, including cell type-specific accessible regions (Figure 2b, S2a-c). We believe this grouping of CREs into “states” captures distinct regulatory programs therefore we call this approach ScregSeg-pi, for <u>p</u>rogram identification, and present its output as the main resource of our study.

Running ScregSeg-pi with 30 states identified states such as 29, 4, 14, 24, 10, 23, 21, 20, which each show clear association with a single cell type (Figure 2b,d-e). State enrichments around ZFIN and sc-RNA-seq marker gene sets (Tables S3,S4) are consistent with those seen for the individual cell type clusters from our one-vs-all differential analysis (Tables S1,S2). On the other hand, states such as 5, 26, 25, 13, and 1, represent accessibility patterns that are specific for multiple cell types (Figure 2b-d), and, therefore, would not be identified by a one-vs-all approach.

We find evidence of multiple distinct regulatory programs being active in a single cell type. For example, cluster 14 is enriched for markers of undifferentiated caudal (somitic) mesoderm and multipotent tailbud precursors with spinal cord, somite, and vascular differentiation potential during body axis extension (Figure S2d)^38–41^. This cluster strongly associates with three states (25, 26, and 13) which likely represent distinct regulatory programs behind the known endothelial, neuronal, and myogenic trajectories for caudal precursors, respectively (Figure S2e, Tables S3-5). Amongst the neuronal clusters, cluster 12 (midbrain) shows strong associations with two states (state 5 and 24). State 5 is associated with all neuronal clusters except cluster 6 (differentiating neurons), and harbors markers of the neurogenesis cascade, while the cluster-specific state 24 includes genomic regions strongly associated with brain spatial identity^42,43^ (Figure S2e, Tables S3-5). This example suggests it is possible to separate regulatory programs driving differentiation of a specific cell lineage (neurogenesis), from the positional segregation of brain regions.

Single-cell accessibility measurements have the potential to shed light on the sequence code of transcriptional regulation. To test this, we utilized convolutional neural networks, which have proven to be powerful tools in DNA sequence analysis^44^, to extract sequence motifs that are associated with individual states of the ScregSeg-pi segmentation model (Figure 3a; see Methods). We find numerous cases where the extracted motifs resemble known motifs whose transcription factors are known to be active in the cell type associated with the state (Figure 3b). These include a MyoD motif detected in several states enriched for muscle specific signal, an ETS motif highly similar to that for FLI1 for the endothelial state 21, a GATA motif from the blood-enriched state 23, and in the neural crest state 20 a motif likely representing the known regulator MITF. These results orthogonally validate our cell type annotations and demonstrate that the ScregSeg segmentation contains sufficient information to train models of sequence regulatory code.

### Bulk assays for chromatin architectures reflect single-cell accessibility dynamics

While accessibility is a highly useful proxy to determine the location of CREs, it does not provide information as to their function or activity. To further characterize the regulatory landscape of 24 hpf embryos, we performed bulk assays for three complementary chromatin modalities: 1) ChIP-seq for five histone post-translational modifications (PTMs) commonly used to define and discriminate promoters, enhancers, and gene bodies (H3K27ac, H3K4me1, H3K4me2, H3K4me3, and H3K36me3), 2) *in situ* Hi-C to detect prominent 3D nuclear organization, and 3) chromatin-associated RNA as a measure of nascent transcription (Figure 4a, S3a). ChIP-seq signal for CRE-associated PTMs served as data to infer the parameters of an HMM for chromatin state annotation^45^ with 11 states (Figure 4b). These states (referred to now as hPTM-states) were further grouped into “promoter-like”, “enhancer-like”, “other”, and “background” types according to overall occurrence (Figure S3b) and spatial patterns around accessible segments proximal and distal to annotated transcription start sites (TSS; Figure S3c,d), which are consistent with previous observations in humans, worms, and flies^45,46^. ScregSeg-fi segments (Figure 1) show enrichment for these different hPTM-states (see Methods) to extents reflecting the type groups (Figure 4c), suggesting current models of CRE architecture apply also to zebrafish. ScregSeg-fi segments were classified into one of the four hPTM-state types (see Methods), confirmed by proximity to annotated TSSs (Figure S3e). Distributions of nascent chromatin RNA-seq counts for segments not overlapping gene bodies confirm the utility of our hPTM-state types since promoter-like states show substantially elevated RNA levels even when located greater than 5 kb from annotated TSSs (Figure S3f).

Previous studies, based on both bulk and single-cell-resolved accessibility measurements^10,11,47^ have observed that promoters show more constitutive accessibility across cell types, whereas enhancers are more dynamic and cell-specific. To describe the cell type-specificity of ScregSeg-fi segments, we calculated the Shannon entropy^11^, for each segment across the 17 identified cell types (Figure 1; see Methods). Indeed, the most constitutive segments show a strong enrichment of promoter-like hPTM-states and proximity to annotated TSSs (Figure 4d, S3g).

Next we integrated hPTM-states and sci-ATAC-seq measurements with 3D genome organization in the nucleus as measured by *in situ* Hi-C^48^ (see Methods). Gene promoters frequently occur at the boundaries of so-called topologically-associated domains (TADs)^49,50^. A common method for determining TAD boundaries from Hi-C data is the insulation score, which measures aggregate interactions that traverse a given genome position^51,52^. Therefore we calculated insulation scores genome-wide and summarized them for each accessible segment. We observed a clear trend for segments with promoter-like hPTM-states (or TSS proximal) to be in more insulated regions and those with enhancer-like hPTM-states (or TSS distal) to be within highly interacting regions (Figure 4e,S3h). Furthermore, constitutively expressed genes are reported to be enriched in TAD border regions^49,53–55^. Therefore we explored the relationship between accessibility-based entropy scores, insulation strength, and hPTM-state, and observed a trend only for more insulated, promoter-like segments to be more constitutively accessible (Figure 4f,S3i).

Co-accessibility of pairs of genomic regions within a certain linear genome distance may be an indication of 3D interactions^9^. To confirm this trend in our dataset, we visualized co-accessibility versus Hi-C interaction scores (Figure 4a, S3a; see Methods) and observed a positive relationship between the two, especially at the high extremes for pairs containing segments with promoter- or enhancer-like hPTM-states (Figure 4g, S3j). At example loci, we clearly observe that strong co-accessibility scores (loops) link both high-scoring 3D interactions (heat map) and regions with high cell type-specific accessibility (Figure 4a, S3a). Furthermore, these co-accessible/interacting regions are assigned to common or related regulatory programs from our ScregSeg-pi analysis (Figure 2), thus showing high concordance between data types and consistency between analysis strategies.

### sci-ATAC-seq reveals cellular changes in *cloche/npas4l* mutants beyond hemato-vascular phenotypes

To test the sensitivity and accuracy of our approach and explore the phenotypic consequences of perturbed gene regulation, we performed sci-ATAC-seq profiling of 24 hpf stage zebrafish embryos with a targeted *npas4l* mutation (*npas4l*^*bns297*^). Such mutants, historically referred to as *cloche*, display distinct alterations of hematopoietic, vascular, and cardiac cell populations^24,25^, therefore we anticipated clear phenotypes in mutant accessibility profiles. After clustering all cells together, homozygous mutants (*npas4l*^*bns297/bns297*^) were separated from their siblings (*npas4l*^*bns297/+*^, *npas4l*^*+/+*^), according to barcodes (Figure 5a). We detect a drastic loss of nuclei within clusters annotated as endothelium and blood (11 and 8, respectively), as reflected by the percentage of mutant nuclei in each UMAP density cluster (Figure 5b). Three additional clusters were less dramatically affected by the mutation: muscle cluster 16 shows an increased mutant nuclei percentage of 50.65% (P-value < 0.0002; Binomial test) compared to the ∼42% average, epidermal cluster 15 shows an increased mutant nuclei percentage of 55.41% (P-value < 2*10^−6^; Binomial test), whereas caudal precursor cell cluster 14 shows a measurable, but non-significant, decrease in mutant nuclei percentage of 34.57% (P-value = 0.05636; Binomial test). Notably, cluster 16 is only populated by nuclei from homozygous mutants and siblings of the *npas4l*^*bns297*^ line (Figure S1i), potentially indicating a heterozygous phenotype.

Our single-cell ATAC-seq dataset allowed us to probe the cell type-specific regulation acting upstream of *npas4l*. Within 30 kb of *npas4l* TSS we observed two cluster-specific accessible regions associated with foreground ScregSeg-pi states 21 and 25 (Figure 2b, Figure 5c). Given their proximity to the *npas4l* TSS, their cell type-specific accessibility in clusters perturbed by *cloche* (cluster 8, 11 and 14), and the absence of long-range interactions with other genomic loci that show cell type-specific accessibility (Figure S4a), we designated these as putative *npas4l* enhancers enh1, enh2a and enh2b. To investigate the potential cell type-specific activity of these putative enhancers, we scanned their underlying genomic sequences against the full JASPAR database^56^ (Figure 5d, S4b-d). In the enh1 sequence we find significant motif matches of blood and mesoderm regulators KLF family, GATA family and LBX2^34,57–60^. Enh2a contains matches for the RFX family motifs^61^, ciliogenesis^62^, and mesodermal regulators (TEAD2^63^, NOTO^64^, EVX2^65^, HOXA5), while enh2b exhibits motif matches for mesodermally expressed transcription factors RXRG and FOXC2^66–68^ and blood regulators GATA1::TAL1. The diverse motif content of these putative enhancer regions underscores the cell type-specificity of their utilization and indicates intricate control of *npas4l* expression.

## Discussion

We present a multimodal resource for the zebrafish community, which integrates sci-ATAC-seq, bulk histone PTMs and Hi-C approaches, to achieve a genome-wide classification of the regulatory architecture determining transcriptional activity in the 24hpf embryo. Using our new tool, ScregSeg, we define regulatory programs specific to one or more of 17 identified cell types and the sequence codes underlying these programs. We find that promoters are mostly constitutively accessible and tend to occur in more insulated 3D neighborhoods and that co-accessible CRE pairs tend to interact in 3D. Sci-ATAC-seq profiling of *npas4l/cloche* mutants revealed unexpected changes in caudal precursor, muscle, and epidermal cell populations. This resource constitutes a solid foundation for future studies in developmental cell biology, systems regulatory genomics, and computational data science, with immediate direct impact on transgenic reporter gene design, candidate identification for perturbation studies, and regulatory sequence labels for further developments in machine learning.

In the wake of advances in sc-ATAC-seq experimental methods, a number of analysis strategies have been developed, which depend on predefined features and/or focus on various aspects such as cell type clustering, motif integration or co-accessibility^9,14,30,69,70^. We developed ScregSeg, a novel HMM segmentation approach for analyzing single cell ATAC-seq data, to address 1) the identification of informative features (e.g. regions with variable accessibility dynamics) for the downstream analysis and 2) the characterization regulatory programs, referred to as ScregSeg-fi and ScregSeg-pi, respectively. We show that ScregSeg-fi generates clearly separated cell type clusters while ScregSeg-pi reveals cluster-specific as well as multi-cell type accessibility profiles in an unsupervised manner (Figure 2), demonstrating the suitability of this approach. This enabled us to dissect several complex regulatory programs potentially representing the neuronal and mesodermal fates in caudal precursors, as well as spatial distribution and lineage progression amongst neuronal clusters. ScregSeg also provides information necessary to identify transcription factor motifs. These biological validations, alongside the scalability to large data, show that ScregSeg is an important new addition to the toolbox of sc-ATAC-seq analysis methods.

Our integration of single-cell datasets with bulk approaches enabled the identification of global trends and multimodal regulatory principles while addressing the issue of bulk signals being dominated by the most prevalent cell types. Enhancers and promoters share many common characteristics and the traditional mark distinguishing them, H3K4me3, may simply reflect higher transcription initiation rates^2,71,72^. Our analyses show H3K4me3-containing hPTM-states and TSS proximity both enrich strongly for constitutive accessibility, thereby suggesting a functional distinction for CREs with this mark. We show that constitutively accessible CREs with promoter-like hPTM-states are associated with highly 3D-insulated regions, refining previous observations that TAD borders are associated with constitutively expressed genes during zebrafish development^55^. Since co-accessible CRE pairs within the same TAD tend to interact in 3D space (Figure 4), our data and analyses raise the potential of assigning a given zebrafish CRE to its target gene if the promoter is not constitutively accessible, as shown previously in a mammalian system^9^.

The novel *cloche/npas4l* phenotypes we observe highlight the power of single-cell methods to quantify small changes in cell numbers during development. The gain in muscle cluster 16 suggests that the lineage commitment of *npas4l* mutant mesodermal precursors is redirected from hemato-vascular to somite-muscle cell types, resembling a phenotype observed upon loss of *etsrp*, a direct target gene of *npas4l*^*73,74*^. As cluster 16 is constituted only by nuclei from homozygous mutant *npas4l* and mixed wildtype/heterozygous siblings (Figure S1i), we propose that heterozygous *npas4l* mutants may exhibit a mild muscle phenotype with a significantly altered chromatin state. However differences in the genetic background or nuclei preparations cannot be ruled out (see Methods). Since the critical role of *cloche* in endothelial cell specification is well established, we hypothesize that the observed decrease in mutant nuclei contribution to cluster 14 represents the known contribution of tailbud precursors to caudal vasculature and its observed differentiation defects in *cloche* mutants^41,75^. Accordingly, ScregSeg-pi identified a regulatory state, state 25, that shows accessibility specific tailbud precursors, pharyngeal mesoderm, and endothelial cells (Figure 2b). An increase in an epidermal population is unprecedented, and will require further investigation.

Finally we leveraged our highly resolved CRE accessibility profiles to explore the *npas4l* locus, where we observe new putative enhancers, enh1, 2a and 2b that exhibit cell type-specificity. The specificity of enh2a/b to caudal precursors supports the importance of *npas4l* in regulating their fate, and we speculate they may regulate *npas4l* early in the hemato-vascular fate decision. Meanwhile the accessibility of enh1 in mature blood/endothelial populations where its RNA levels are not detectable could support a purported negative-feedback regulation of *npas4l’*s transient expression^25^. Exploration of these enhancers is one example of how the data generated from this study provides an extensive resource for follow-up studies.

## Methods

### Embryo preparation for sci-ATAC-seq

For wild type experiments the AB/TL strain was used. All zebrafish maintenance and procedures were conducted in accordance with standard laboratory conditions and animal procedures approved by the local authorities (LAGeSo, Berlin, Germany). Timed matings were set up between AB/TL adults and embryos were maintained at 28.5°C for 24 hpf from the time of fertilization. Staging and consistency within the clutch was confirmed by morphological criteria^27^.

Chorions were removed by incubating in 15 ml pronase E at 1 mg/ml for 10 min with continuous shaking. Pronase was removed by five washes with 200 ml egg water (60 µg/ml Ocean salt (Red Sea), 3 µM Methylene blue). For the first two wildtype experiments, embryo yolks were removed by placing 100 embryos in 500 µl de-yolking buffer (55mM NaCl, 1.8mM KCl, 1.25mM NaHCO_3_) and pipetting 10 times with a P100 pipette. Embryos were left to sink to the bottom, then de-yolking buffer was removed and 5 washes with egg water were performed. Batches of 25-50 embryos were distributed into 1.5ml eppendorf tubes, egg water removed and snap frozen in liquid nitrogen and maintain at -80°C.

Embryos with mutated *npas4l* alleles were obtained from intercrosses of *npas4l*^*bns297*^ heterozygotes in a *Tg(fli1a:nls-GFP)*^*y7*^ background ^74^. Homozygous *npas4l* mutant (*npas4l*^*bns297/bns297*^) embryos were separated from heterozygous and homozygous wild type siblings (*npas4l*^*bns297/+*^, *npas4l*^*+/+*^) based on the loss of fli1a-GFP+ endothelial cells 24 hours after fertilization. Chorions were removed but yolks were left intact. Embryos were flash frozen in liquid nitrogen, stored at -80°C and transported on dry ice.

### Nuclei preparation from zebrafish embryos for sci-ATAC-seq

Embryos were thawed in 2 ml of cold lysis buffer (CLB) (10 mM Tris-HCL, pH 7.5, 10 mM NaCl, 3 mM MgCl_2_, 0.1% IGEPAL CA-630)^7^ supplemented with protease inhibitors (Complete Protease Inhibitor Cocktail, EDTA-free, Roche). Embryos were homogenized in a Dounce homogenizer then incubated in cold lysis buffer at 4°C for 1 hour. Nuclei were then strained through 35 micron strainer caps of Corning Falcon test tubes (Thermo Fisher Scientific).

### Nuclei preparation from sea urchin embryos for sci-ATAC-seq

Sea urchin embryos were S. purpuratus, wild caught in Monterey, California, USA. Embryos obtained 30-48 hpf were fixed for 30 min in 5 mM DSP then quenched with 20 mM Tris pH 7.4 and stored at 4°C. For nuclei preparation, fixed embryos were thawed in 10 ml HB buffer (15 mM Tris, pH 7.4, sucrose 0.34 M, NaCl 15 mM, KCl 60 mM, EDTA 0.2 mM, EDTA 0.2 mM) and then homogenized in a 15 ml Dounce homogenizer 20x with a loose pestle, and 10x with a tight pestle. The homogenate was filtered through Miracloth (Merck Millipore) and rinsed with HB buffer, followed by centrifugation at 3500 g for 5 min at 4°C, discarding the supernatant, twice. The pelleted nuclei were resuspended in cold PBS with 0.1% Triton X-100 and filtered through a 20 µM Nitex membrane, then spun down and resuspended in 1ml CLB.

### Tn5 transposome preparation for sci-ATAC-seq

Tn5 was generated by the MDC Protein Production & Characterization Platform from Addgene plasmid #60240 according to^76^ at 1.95 mg/ml with the following minor modifications: buffers lacking Triton X-100 were used for the chitin column and dialysis, and final storage was in 50 mM Hepes-KOH pH 7.2, 0.8 M NaCl, 55 % Glycerin, 0.1 mM EDTA, 1 mM DTT. For each experiment 96 uniquely indexed transposon complexes were generated according to^77^ with minor adaptations. Firstly, twenty uniquely indexed transposons were made by annealing a uniquely indexed oligonucleotide (Sigma-Aldrich) containing a Tn5 mosaic end sequence at its 3’ end, to a complementary universal 5’-phosphorylated 19 bp mosaic end oligonucleotide. Oligonucleotides were mixed in a 1:1 molar ratio, giving a final concentration of 100 µm under the thermocycling conditions: 95°C for 5 minutes, cool to 65°C decreasing 0.1°C/second, 65°C for 5 minutes, cool to 4°C decreasing 0.1°C/second. Each annealed oligonucleotide transposon was mixed with Tn5 (1.95 mg/ml) at a ratio of 0.143:1 and incubated for one hour at 25°C. Of the 20 indexed oligonucleotides, 8 contained an adaptor that could be bound by indexed Illumina P5 primers (i5 oligonucleotides) and 12 contained an adaptor that could be bound by indexed Illumina P5 primers (oligonucleotide sequences were obtained from supplementary information of ^14^). To make 96 unique transposome complexes each i5 transposome could be mixed with each i7 transposome at a 1:1 ratio in columns 1-12 and rows A-H of a 96 well plate. Transposome complexes were stored at -20°C.

### sci-ATAC-seq method

Our protocol for generating sci-ATAC-seq data was largely following^10,14^, with some modifications. Purified nuclei were stained with DAPI (4 µM) and 2500 were sorted into each well of a 96 well plate containing 19 µl of tagmentation buffer (10mM TAPS-NaOH, pH 8.8, 5 mM McCl_2_, 10% DMF, 6.6 mM Tris-HCl, 6.6 mM, 0.066 % IGEPAL CA-630) using a BD FACS Aria III (BD Biosciences). For tagmentation, 1 µl of uniquely barcoded Tn5 transposome was added to each well of the 96 well plate containing tagmentation buffer and nuclei. Plates were spun for 30 seconds at 500 x g and then incubated for 30 minutes at 37°C. Following tagmentation, 40 µl of 40 mM EDTA supplemented with 0.3 mM spermidine was added to each well and the plate incubated for 15 minutes at 37°C. Nuclei and buffer from all wells were pooled in a reagent reservoir and passed through a 35 micron strainer into Corning Falcon test tubes (Thermo Fisher Scientific). DAPI was added and nuclei were sorted again with a BD FACS Aria III. For the second sort, 25 nuclei were sorted into each well of 96-well plates (8-10 plates per experiment) containing 12 µl of nuclear lysis buffer (11 µl of EB buffer (Qiagen) supplemented with 0.5 µl of 100X BSA and 0.5 µl of 1% SDS). The 96-well plates from the second sort were stored at -20°C until ready for PCR amplification.

Before PCR amplification, each plate was incubated at 55°C for 15 minutes then 1 µl of 12.5% triton-X100 added per well to quench the SDS. To each well a unique combination of indexed P5 and P7 PCR primers^14^ was added (0.5 µM final concentration each), 10 µl of NEBNext Ultra II Q5 Master Mix (NEB) then immediately amplified in a thermocycler under the conditions: 72°C for 3 minutes, 98°C for 30 seconds, 18 cycles: 98°C for 10 seconds, 63°C for 30 seconds, 72°C for 1 minute, hold at 4°C. Before amplifying a whole plate, the number of cycles was determined from several test wells that were sorted into a separate plate and monitored by qPCR with the addition of SYBR green to the PCR mix. In all experiments here 18 cycles were used.

After PCR amplification, 96-wells of each plate were pooled, cleaned up with DNA Clean & Concentrator-5 columns (Zymo) and then large fragments (above 1000 bp) removed with 1X AMPure beads (Beckman Coulter). The concentration of libraries was measured with Qubit dsDNA HS Assay (Thermo Fisher Scientific) and quality checked with Bioanalyzer DNA High Sensitivity Kit (Agilent).

### Sequencing of sci-ATAC libraries

Equimolar libraries from each 96 well plate were pools and sequenced with NextSeq500 (Illumina) High Output, 2 × 150 bp loading at a concentration of 1.6 pM. Custom primers^14^ and a custom sequencing recipe^77^ were used to sequence the following read lengths: 110 bp + 45 bp + 110 bp + 39 bp (Read 1 + Index 1 + Read 2 + Index 2).

### Sci-ATAC-seq Analysis

#### Preprocessing

The raw sequences were trimmed using flexbar with parameters ‘-u 10 --min-read-length 50’ using the adapter sequence ‘CTGTCTCTTATACACATCTG’. Reads were mapped against danRer11 using ‘bowtie2 -X 2000 --no-mixed --no-discordant --very-sensitive’. Chromosomes whose names contain the patterns chrM, _random and chrUn were removed from the analysis and only reads with mapping quality of at least 10 were retained. We corrected sequencing errors in barcodes by mapping the sequenced barcodes against the reference barcode universe using bowtie2 with default parameters. Only barcodes with a mapping quality of at least 5 and no more than two mismatches with the reference barcodes were retained. Finally, reads were deduplicated within each barcode using a custom script. A subset of the barcode universe was used to sequence Zebrafish and Sea urchin simultaneously in order to detect barcode collision events. The barcode collision rate was estimated as described previously using the Birthday paradox^8^.

#### Defining the regions of interest using ScregSeg-fi

We binned the genome in 1 kb regions and constructed a single-cell count matrix with dimensions number of cells by number of genome-wide bins. Fragments were counted at the midpoint and each entry of the matrix was trimmed to be at most four to mitigate the influence artifacts. To exclude poor quality cells, we only retained barcodes with at least 1000 and at most 30000 fragments, leading to 21136 barcodes used at this stage.

We developed a hidden Markov model (HMM), called ScregSeg, to segment the single cell ATAC-seq profiles along the genome. The HMM uses Dirichlet-Multinomial emission probabilities, one associated with each state, to describe the cross-cell accessibility profile of each bin in the genome. This allows to associate potentially cell-type-specific accessibility profiles with distinct states.

We utilized an HMM with 50 states and fitted the model using the Baum-Welch algorithm for 300 iterations starting from random initial parameters. Since the model might get trapped in a poor local optimum, we restarted the training process seven times using different random initial weights and eventually used the model that obtains the best overall log-likelihood score. However, in general, the solutions found with different random initial weights were relatively comparable across runs.

We performed state calling using the Viterbi algorithm and only retained rare/informative states which cover at most 1.5% of the genome each. States that cover a larger fraction of the genome are considered background states. Furthermore, we used only state calls with posterior decoding probability of at least 0.9 to eliminate uncertain state calls and merged bookended bins if they belonged to the same state. The set of intervals obtained by this approach constitute the regions of interest, which were used for the downstream analysis (e.g. Latent Dirichlet Allocation).

#### Dimensionality reduction and clustering

We constructed a single-cell count matrix using all cells across all samples and using the regions of interest defined by ScregSeg-fi (see above). The count matrix was subjected to filtering requiring at least one fragment per region across cells and at least 200 fragments per cell across the ROI regions which led to 23008 barcodes being used for the remaining analysis. We fitted a Latent Dirichlet Allocation model using the cisTopic package using 30 Topics, collapsed Gibbs sampling, a burn-in of 500 and 1000 sampling iterations^30^. The resulting cell-topic matrix was z score-normalized and sample specific batch effects were corrected for by regressing out the sample labels with a linear regression model. The cell-topic matrix was further used to construct a 2D UMAP. We performed density clustering on the UMAP to group together cells in distinct subpopulations. We created pseudo-bulk signal tracks based on cells within each density cluster. Given some query regions (e.g. known marker genes), cell-specific enrichment scores were determined using the AUCell score provided by cisTopic^30^. These enrichment scores were used to highlight marker accessibility in the UMAP.

#### Differential peak calling

Cluster-specific marker regions were determined by performing one-vs-all differential accessibility analyses using DESeq2 for each density cluster in turn using the regions of interest identified by ScregSeg-fi.

For each cluster, regions with a minimum log2-fold change of one and an adjusted p-value of at most 10% were reported as cluster-specific regions. In addition, the top 500 regions with respect to the log-fold change were reported regardless of the above constraints.

#### Extracting ZFIN-derived annotation features

We compiled body-part specific gene sets using annotation from the ZFIN database ^31^. To this end, we downloaded the gene-body-part association and extracted body-parts present in the 24 hpf developmental stage. We only used annotation data for the publication ids ZDB-PUB-040907-1, ZDB-PUB-051025-1 or ZDB-PUB-010810-1 to ensure consistent quality and remove body parts with less than 6 genes.

#### Extracting scRNA-seq-derived marker genes

We compiled gene sets based on cluster-specific genes for published single-cell RNA-seq data^23^. To this end, we downloaded the single-cell RNA-seq count matrix along with the cell clustering information from Wagner et al 2018. We employed scVI^78^ to determine one-vs-all differential gene expression for each cluster in turn based on the 24 hpf single-cell data. We use the top 20 most differential genes per cluster to constitute the scRNA-seq-cluster gene set.

#### Gene enrichment analysis per cluster

For each density-cluster, we determined whether the differentially accessible regions are significantly enriched around the gene sets defined by ZFIN and scRNA-seq data using a hypergeometric test. To this end, we mapped each differentially accessible region to the nearest gene and ensured that each gene was counted only once, in case of multiple marker regions mapping to the same gene.

#### Genome segmentation for identifying regulatory programs - ScregSeg-pi

We binned the genome in 500 bp regions. Then we constructed a count matrix containing pseudo-bulk fragment counts across the genome resulting in a count matrix with size (number of bins) x (number of density clusters).

We applied the ScregSeg segmentation model with Dirichlet-Multinomial emission probabilities for segmenting the genome similarly as described above.

We fitted a model with 30 states for 100 iterations using the Baum-Welch algorithm. Starting the fitting process from different initial weights produced similar results. Nevertheless, we repeated model fitting seven times with different random initial weights to minimize the chance of obtaining a poor local optimum. Finally the model with the best log-likelihood score was selected.

To visualize the state-cluster association, we denote c and s as the cluster and state indices and computed the log-likelihood ratio 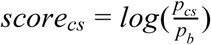 using the state-specific coverage profile 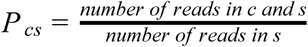 and the background coverage profile 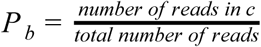.

#### Feature enrichment score per states

In order to assign biological function to states, we developed a statistical test based on the abundance of state calls around the TSS of genes in the gene sets. To this end, we counted the number of state calls *o*_*i*_ for each state *i* across the region defined by the gene set. The expected number of state counts in a region of the same size N was computed *e*_*i*_ = *Np*_*i*_ where *p*_*i*_ denotes the stationary probability of state i of the segmentation model. We used the enrichment score: 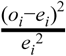 if (*o*_*i*_ − *e*_*i*_) > 0. Otherwise, the score is zero. The feature enrichment scores were used to associate functional gene sets with states and for visualization purposes. As gene sets we use the ZFIN and scRNA-seq extracted marker genes as described above and we expanded the regions around the TSSs by +/- 10k.

#### Query marker genes for a given state

In order to extract marker genes, we utilized the log ratio between the proportion of observed state calls covering the gene body +/-10 kb and the proportion of state calls expected by chance in a region of the same size based on the stationary distribution of the HMM. A high positive score indicates an excess of state calls for a particular state relative to its genome-wide state abundance.

### Testing cell count differences between *npas4l* mutants and siblings

This test was performed on sci-ATAC-seq profiles from *npas4l* mutant embryos (*npas4l*^*bns297/bns297*^) and their siblings (*npas4l*^*bns297/+*^, *npas4l*^*+/+*^). *npas4l* mutant cells were separated from siblings based on their tagmentation barcodes. We tested for enrichment or depletion of cells from *npas4l* mutants vs siblings using a binomial test for each density cluster. The success probability was determined by (total number of *npas4l* mutant cells) / (total number of *npas4l* mutant and sibling cells).

#### Motif discovery using neural networks

We utilized convolutional neural networks to predict the state probability from the segmentation model weighted by the read counts from the underlying DNA sequence and thereby extracting associated motifs. To this end, we introduce the target score for the regression task as *s*_*ij*_ = *d*_*ij*_ × *r*_*i*_ for region i and state j where *d*_*ij*_ denotes the posterior decoding probability of region i and state j and *r*_*i*_ denotes the aggregated read counts for region i. That is, the score captures the cross-cluster accessibility pattern associated with a state while also emphasizing regions with high read counts. For each state, we extracted the top 100k regions according to that score of which we kept the top 15k and the bottom 15k sequences for model training and evaluation. These can be considered as positive and negative sets. As input to the convolutional neural network we extracted the 500 bp DNA sequences associated with the training and evaluation regions extended by +/- 250 bp and converted them to one hot encoding. The network uses a convolutional layer with 100 kernels, 13 bp kernel length and sigmoid activation to scan both strands of the DNA sequence. Subsequently, the maximum activation across the strands is propagated forward and subjected to global max pooling, dropout with 50% and a linear output node. We choose the sigmoid activation in the initial layer due to its relationship with representing Bernoulli random variables, which, after normalization, allows us to approximately interpret the kernel weights as log-likelihood ratios and thus position weight matrices.

The network was trained on all regions, except for regions on chromosome 1 and 5 which were used as validation set. Training was performed using ADAM by minimizing the mean absolute error for at most 300 epochs with batch size 32 and early stopping with a patience of 20 iterations. After model fitting, the 10 kernels whose maximum hidden activations per sequence after the first convolution layer individually correlated most with the state-specific score were extracted, normalized to represent PWMs and reported as de novo motifs. These motifs were matched against motifs from JASPAR 2018^56^, non-redundant vertebrates using TOMTOM^79^.

### ChIP-seq

Embryos were collected 24 hpf and fixed in 0.5% formaldehyde (Carl Roth #4979.1) for 15 minutes as previously published for *D. melanogaster*^*80*^ with minor modifications: heptane was not added to the fixation buffer since unlike *D. melanogaster* zebrafish does not have a cuticula. Nuclei were extracted according to ^81^ and sonicated for 16 cycles (30s ON, 30s OFF, on high setting) in a Bioruptor Plus (Diagenode) to achieve a DNA fragment size below 500 bp. ChIP was performed using True MicroChIP Kit (Diagenode #C01010130) according to the manufacturer’s instructions with the following modifications: primary antibody was incubated at 4°C overnight and the reverse crosslinking was done overnight. The following antibodies were used: H3K4me1 (abcam #ab8895), H3K4me2 (abcam #ab32356), H3K4me3 (abcam #8580), H3K27ac (abcam #ab4729), H3K36me3 (abcam #ab9050). The library was prepared using NEXTflex qRNA-Seq Kit v2 (BioScientific #5130-12, discontinued) according to the instructions for qChIP-Seq and paired-end sequencing (2×75nt) was performed on a NextSeq 500/550 using a HighOutput v2 Kit for 150 cycles (Illumina #FC-404-2002, discontinued).

### ChIP-seq processing

UMIs were extracted from paired-end reads using UMI-tools^82^ and mapped to the danRer11 genome assembly using Bowtie2^83^; -X 2000 --no-discordant --no-mixed). Mapped reads were filtered for MAPQ 30 and deduplicated using UMI-tools. Input-subtracted .bigwig for visualization (--operation subtract --binSize 50 --scaleFactorsMethod None --normalizeUsing CPM --smoothLength 250 --extendReads) and .bedgraph for HMM (--operation subtract --binSize 1 --scaleFactorsMethod None --normalizeUsing CPM --extendReads; see below) tracks were generated using deepTools^84^. Reads were converted to bedpe files using bedtools^85^. Peaks were called using JAMM^86^ considering both replicates separately (-r window -e 1 -b 250 -t paired).

### Histone PTM HMM

Signals and peak calls from histone PTM ChIP-seq data were used as input for generating a HMM segmentation model as previously described^45,72^. Briefly, using bedtools intersect and map^85^, genome-wide 10 bp resolution tracks were generated for each factor such that places where peaks were called were assigned values from ChIP-seq signal files and where no peaks were called assigned values of zero. These signal tracks for chromosomes 1 and 2 were then used as input to bw.r (https://github.com/mahmoudibrahim/hmmForChromatin) to learn the model and the resulting model used to decode the rest of the genome states using decoding.r (https://github.com/mahmoudibrahim/hmmForChromatin). The model was learned with increasing number of states until patterns of state coverage around segments proximal to annotated TSSs resembled previously observed patterns across metazoans^46^, leading to the selection of a model with 11 states plus one background state where no ChIP-seq peaks were called and/or ChIP-seq signals were <= 0.

To classify sci-ATAC-seq regions as being enriched for a given histone PTM state, we developed a score where the state coverage for a given region (obtained using bedtools annotate) is divided by the sum of the observed state coverage for the region set, and then took the log of the ratio between this normalized coverage and the expected coverage of the state for that region size given the genome-wide state probabilities. The region was classified as the state with the highest score. For Figure 4c, we applied this classification to all genome-wide 1 kb bins, counted the number of classifications for each state, and divided that number by the total number of 1 kb bins to get expected state classification fractions. We then applied the classification to the unmerged sci-ATAC-seq foreground 1 kb bins, calculated the observed classification fractions for each state, and plotted the log2 ratio between this number and the expected state classification fractions.

### Cellular fractionation

Embryos were collected 24 hpf, dechorionated and homogenized on ice in buffer N (10 mM HEPES pH 7.5, 250 mM sucrose, 50 mM NaCl, 5 mM MgCl_2_, 1 mM DTT, 1X Complete Protease Inhibitor (Roche #11697498001) and 20 U/ml SUPERase-In RNase Inhibitor (Thermo Fisher Scientific #AM2696) using a Dounce homogenizer. After allowing the debris to settle for 5 minutes on ice the supernatant was then washed in PBS, loaded on a sucrose cushion, centrifuged at 1000 g at 4°C and further fractionated according to^87^. RNA was extracted using Trizol (Thermo Fisher Scientific #15596018) and Direct-zol RNA MiniPrep Kit (Zymo Research #R2052) according to the manufacturer’s instructions. The library was prepared using NEXTflex Rapid Directional qRNA-Seq Kit (BiooScientific #NOVA-5130-01D) according to the manufacturer’s instructions and paired-end sequencing (2 × 75 bp) was performed on a NextSeq 500/550 using a HighOutput v2 Kit for 150 cycles (Illumina #FC-404-2002, discontinued).

### Chromatin RNA-seq processing and analysis

Unique molecular identifiers (UMIs) were extracted from .fastq files using UMI-tools^82^ and reads trimmed using fastx_trimmer from the FASTX-toolkit (http://hannonlab.cshl.edu/fastx_toolkit/). Reads were then filtered for ERCC spike-in reads and rRNA by mapping to a custom index with Bowtie 1^88^. Trimmed, filtered, reads were then mapped using STAR^89^. Mapped .bam files were then subjected to PCR deduplication using UMI-tools^82^, followed by conversion to .fastq and remapping with STAR to generate final mapped files and normalized coverage tracks. For sci-ATAC-seq segment chromatin RNA quantification, coverage tracks were created for each genome strand using deepTools^84^ and then summed within a 5 kb window centered around the segment midpoint using bedtools map^85^.

### sci-ATAC-seq Entropy

Foreground sci-ATAC-seq segments were counted for reads from density cluster-collapsed .bam files using bedtools multicov. A pseudocount of 1 was added to the matrix before per-cluster depth normalization. Then the Shannon entropy was calculated for each segment’s normalized count vector across the clusters using the following equation:

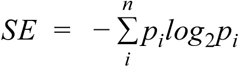

Where *p* is the probability of ATAC-seq signal in cluster *i* for a given segment and *n* is all the sci-ATAC-seq clusters.

### Co-accessibility

Foreground sci-ATAC-seq segments were measured for co-accessibility using Cicero^9^ with cisTopic topic probabilities and topic-based UMAP coordinates (see above Methods “Dimensionality reduction and clustering”) as reduced dimension information, but otherwise with default parameters.

### *In situ* Hi-C

Embryos were collected 24 hpf, dechorionated, fixed in 1% formaldehyde in PBS, quenched and washed as in^80^. Nuclei were extracted according to^81^ using the cell lysis buffer (10 mM Tris–HCl pH 7.5, 10 mM NaCl, 0.5% IGEPAL (CA-630, Sigma, I8896) and a Dounce homogenizer.

Hi-C library preparation was performed as previously described^48^ with modifications. In brief, 25 × 10^6^ isolated nuclei were divided in 5 aliquots and digested overnight with 1500U of HindIII per aliquot. After biotin-fill in, proximity ligation was carried out in each aliquot with 100 units of T4 DNA ligase (Invitrogen) at 16°C overnight. DNA was purified by reverse crosslinking and DNA precipitation, and biotinylated nucleotides were removed from unligated fragments ends with 1U T4 DNA polymerase per µg DNA for 4 hours at 12°C. DNA was sonicated to 300 – 500 bp, using the Bioruptor Plus and size selected for fragments between 300 and 500 bp with AMPure XP beads (Beckman Coulter). Biotinylated DNA fragments were pulled down with MyOne Streptavidin C1 beads (Invitrogen), end-repaired, A-tailed, and TruSeq sequencing adapters were ligated to the DNA fragments with 15U T4 DNA ligase (Invitrogen) overnight at 16°C, shaking at 750 rpm. Adapter-ligated DNA was amplified for 6-8 cycles using Herc II Fusion DNA polymerase (Agilent). PCR products were purified with AMPure XP beads and subjected to Illumina paired-end sequencing (2x 75 bp).

### *In situ* Hi-C analysis

*In situ* Hi-C reads were initially processed using the Juicer pipeline^90^. TADs were called using the insulation method^51^ with default settings (--is500000--nt0--ids250000--ss0--immean) after dumping valid interactions at 25 kb binned resolution using juicer-tools and converting formats using HiTC ^91^. For scoring Hi-C interactions, valid interactions were dumped by juicer-tools at fragment resolution, filtered to remove interactions less than 20 kb apart and format converted using custom scripts, and then subjected to shuffling and scoring using the SHAMAN method^92^.

Insulation scores for sci-ATAC-seq segments were calculated as the mean signal from 40 kb windows surrounding the midpoints of the segments using bedtools slop and map^85^.

For Hi-C/co-accessibility analysis, foreground sci-ATAC-seq segments used for clustering were measured for co-accessibility (see above). Segment pairs were then filtered to remove pairs less than 25 kb (custom scripts) apart and to be within the same TADs using bedtools pairtobed^85^. Filtered segment pairs were then assigned a SHAMAN interaction score^92^, using a slightly modified version of “shaman_generate_feature_grid” (https://bitbucket.org/tanaylab/shaman/src/default/) in which pair relationships are pre-determined and not re-calculated. For all filtered segment pairs, the maximum SHAMAN score was retrieved within a 5 kb grid around the interaction point from the segment midpoints.

### Plotting

All plots for Figure 4 and S3 were generated using ggplot2^93^, except for the Hi-C heatmaps which were plotted using “shaman_plot_map_score_with_annotations” (https://bitbucket.org/tanaylab/shaman/src/default/), the browser shots which were plotted using CoolBox (https://github.com/GangCaoLab/CoolBox), the co-accessibility loops which were plotted using the built-in Cicero plotting function^9^, the histone PTM state coverage around annotated segments which was plotted from built-in R plot function, and the histone PTM state/mark enrichment heat map which was plotted by default from the HMM scripts (https://github.com/mahmoudibrahim/hmmForChromatin).

### Motif scanning in putative enhancers of *npas4l*

Sequences form putative enhances were obtained from their genomic coordinates using bedtools getfasta v2.27.1^85^ with reference genome danRer11 described above. A 0-order Markov local background was generated for each putative enhancer from its sequence plus its 250 bp flanking sequences using fasta-get-markov from the MEME Suite 4.11.3^79^. Each putative enhancer sequence was then scanned for matches to motifs from the JASPAR vertebrate database^56^ using FIMO from the MEME Suite 4.11.3^94^ with the aforementioned background.

## Acknowledgements

We thank Anja Schütz and the team of the MDC Protein Production & Characterization Platform for Tn5 transposase protein production; Dr. Hans-Peter Rahn, Caroline Braeuning and the team of MDC Flow Cytometry Technology Platform for technical support; Ronny Schäfer, Jana Richter, Robby Fechner, Angelica Ospina and the team of MDC Zebrafish Facility for zebrafish maintenance and technical support. We extend our thanks to Pedro Olivares Chauvet for advice on Hi-C analysis, Alex Glahs for advice on nuclei extraction and initial FACS support, Rebecca Worsley Hunt for advice on ChIP-seq processing, Martin Burkert and Dermot Harnett for coding assistance, and Luca Tosti for critical scientific discussion. This research was funded by a Helmholtz Association networking grant “From Sparse to Big Data: data imputation & data fusion for massive sparse data. WK received support from the German Federal Ministry of Education and Research (de.NBI; FKZ 031L0101B). SAL is supported by the Stiftung Charite as a BIH Delbrück Fellow.

## Author Contributions

ACM established and performed sci-ATAC-seq experiments with advice and assistance from DV, SAL, AH, IB, DAG and AT and support from JPJ, as well as various downstream sci-ATAC-seq bioinformatics analyses. WK developed the sci-ATAC-seq processing pipeline and ScregSeg and performed most of the sci-ATAC-seq computational analyses with support from AA and UO. DV performed ChIP-seq and chromatin RNA-seq with assistance from AH and with zebrafish support from AMM and DP, as well as Hi-C experiments together with RK supported by AP, and assisted with ChIP-seq and chromatin RNA-seq data processing. SAL performed ChIP-seq, chromatin RNA-seq, and Hi-C analyses. KM phenotyped and prepared *cloche* embryos with support from DYRS. ACM, WK, and DV made the figures. ACM, WK, and SAL wrote the text. DV and UO edited the text. RK, KM, DP, JPJ, AA, AP, DAG, and DYRS gave comments and suggestions on the manuscript and performed minor editing. SAL and UO designed the study.

## Supplementary figures

**Supplementary figure S1.**
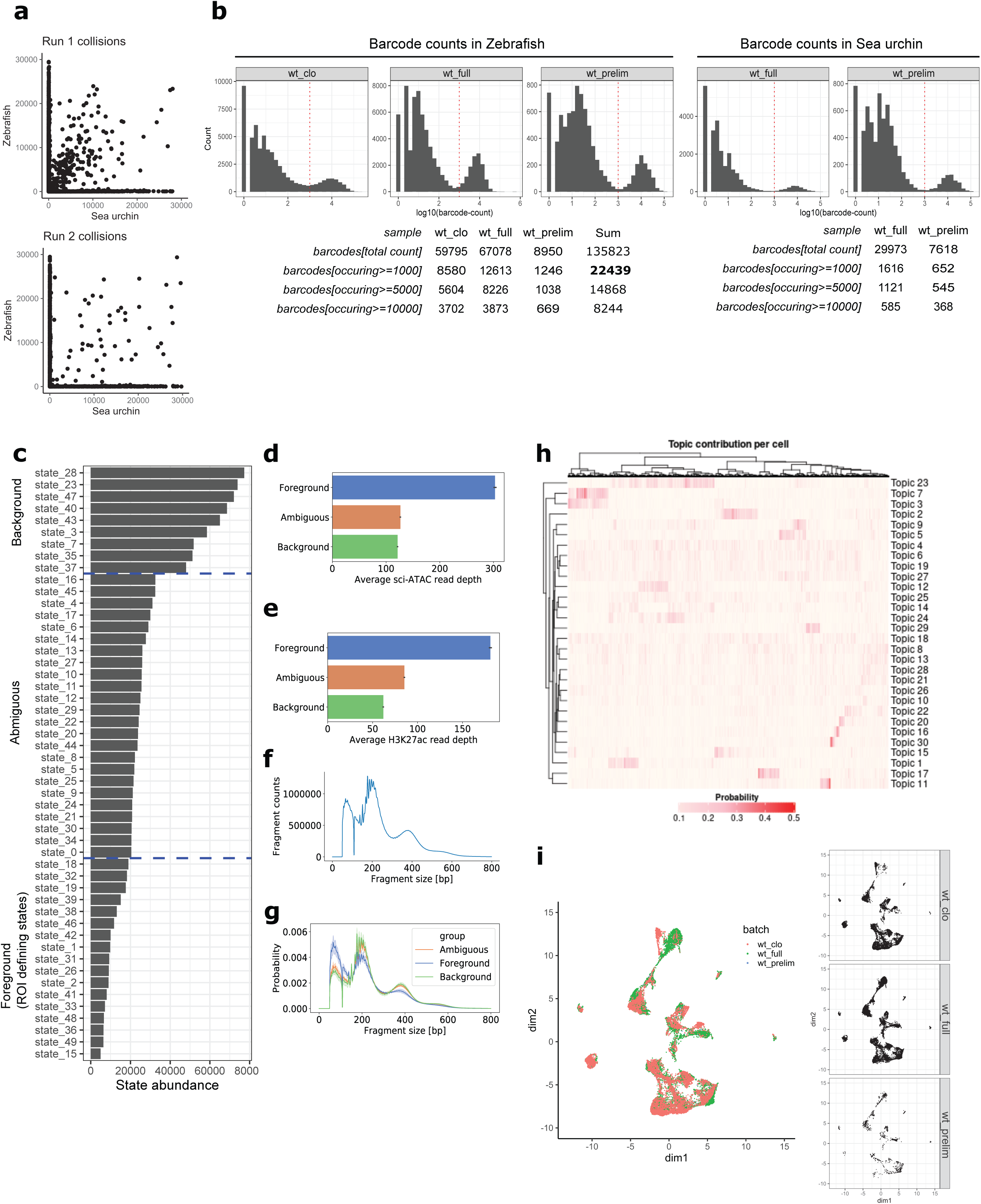
a. Assessment of the barcode collision events. Number of uniquely mapped reads per barcode that align to zebrafish (danrer11) and sea urchin (spur3.1) in the first and second run. Only barcodes with at least 1000 fragments in either species are considered. b. Histogram of the number of uniquely mapped reads per barcode from alignment to zebrafish (danrer11) and sea urchin (spur3.1) in the two wild-type runs (“wt_full”, “wt_prelim”) and only zebrafish (danrer11) in the *npas4l*^*bns297*^ cloche mutant (“wt_clo”). c. State abundance of the ScregSeg-fi model reflects how biologically informative states may be. Rare states tend to reflect cell-type specific accessibility patterns, whereas abundant states describe the background accessibility profile. d. Rare states tend to exhibit higher read counts aggregated across cells. e. Rare states tend to exhibit higher enhancer signals (as measured by bulk H3K27ac ChIP-seq), suggesting that they capture relevant regulatory accessibility signals. f. Overall fragment size distribution. g. Rare states show a distinct fragment size distribution profile compared to background states. In particular, rare states tend to be enriched for short size fragments, which are associated with nucleosome-free regions. h. Heatmap representing the probability of each topic’s contribution per cell. i. UMAP representation of all filtered cell types after batch correction, colored by batch. Batches are defined as the three sci-ATAC-seq experiment runs: two runs with wild-type embryos (“wt_full”, “wt_prelim”) and one run with *npas4l*^*bns297/+*^, *npas4l*^*+/+*^, *npas4l*^*bns297/bns297*^ embryos (“wt_clo”).

**Supplementary figure S2.**
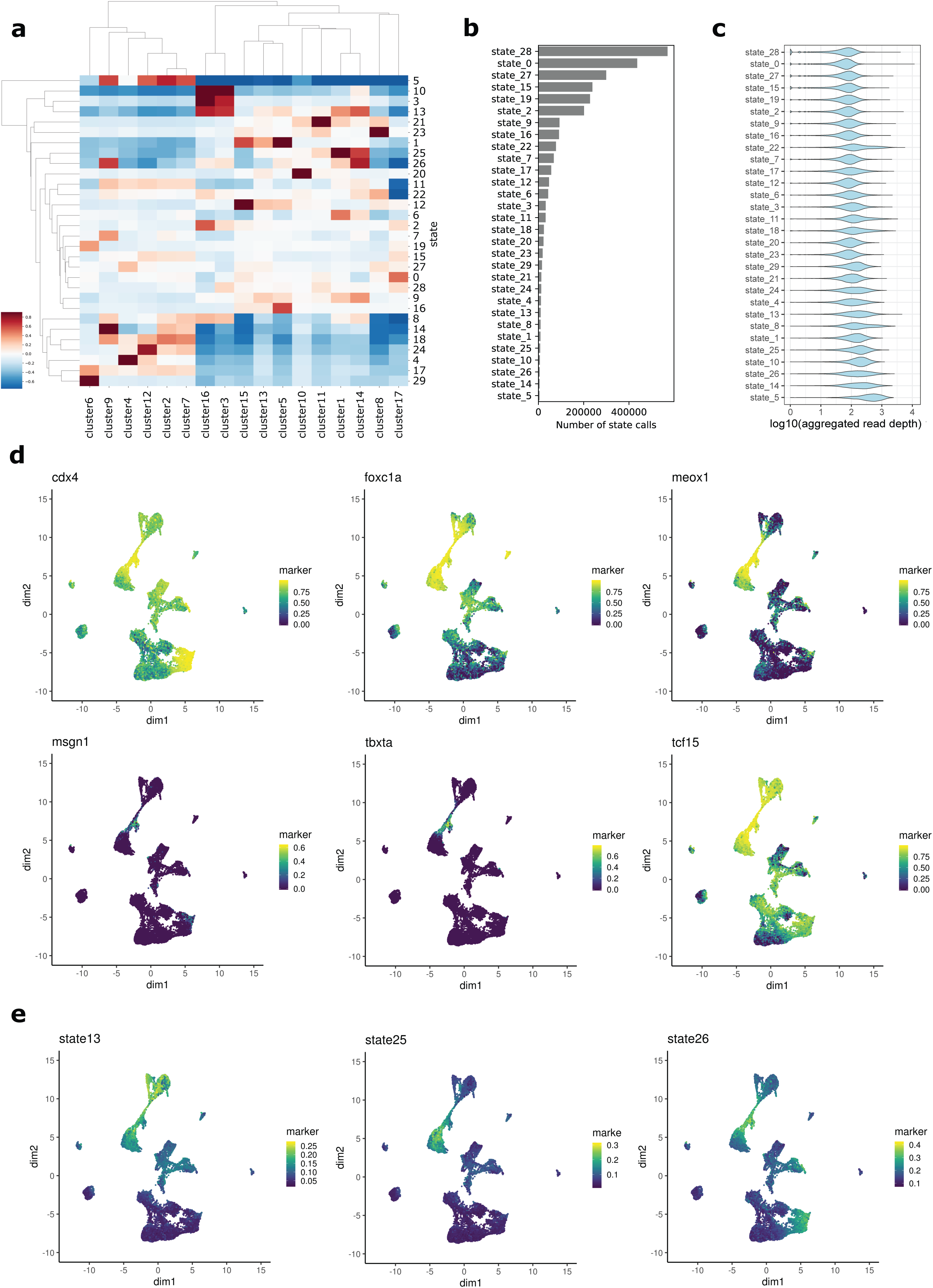
a. Heat map representing the association between each state with each cluster based on the log-fold enrichment between the ScregSeg-pi states’ emission probabilities and the genomic background coverage profile (see Methods). b. State abundance of the ScregSeg-pi model. c. Aggregated read depth (across all clusters) associated with the states. d. Per cell distribution of accessibility at promoters of marker genes of the putative constituent cells of cluster14: neural mesodermal progenitors (*tbxta, cdx4*) and presomitic mesoderm (*foxc1a, meox1, msgn1, tcf15*), represented in UMAP space. Color represents the rank-based AUCell enrichment score for a given region^30,95^. e. Per cell distribution of accessibility at the gene body of genes mapping to segments with the top 100 logFC enrichment in states 13, 25 and 26 represented in UMAP space. Coloring of enrichment as in D).

**Supplementary figure S3.**
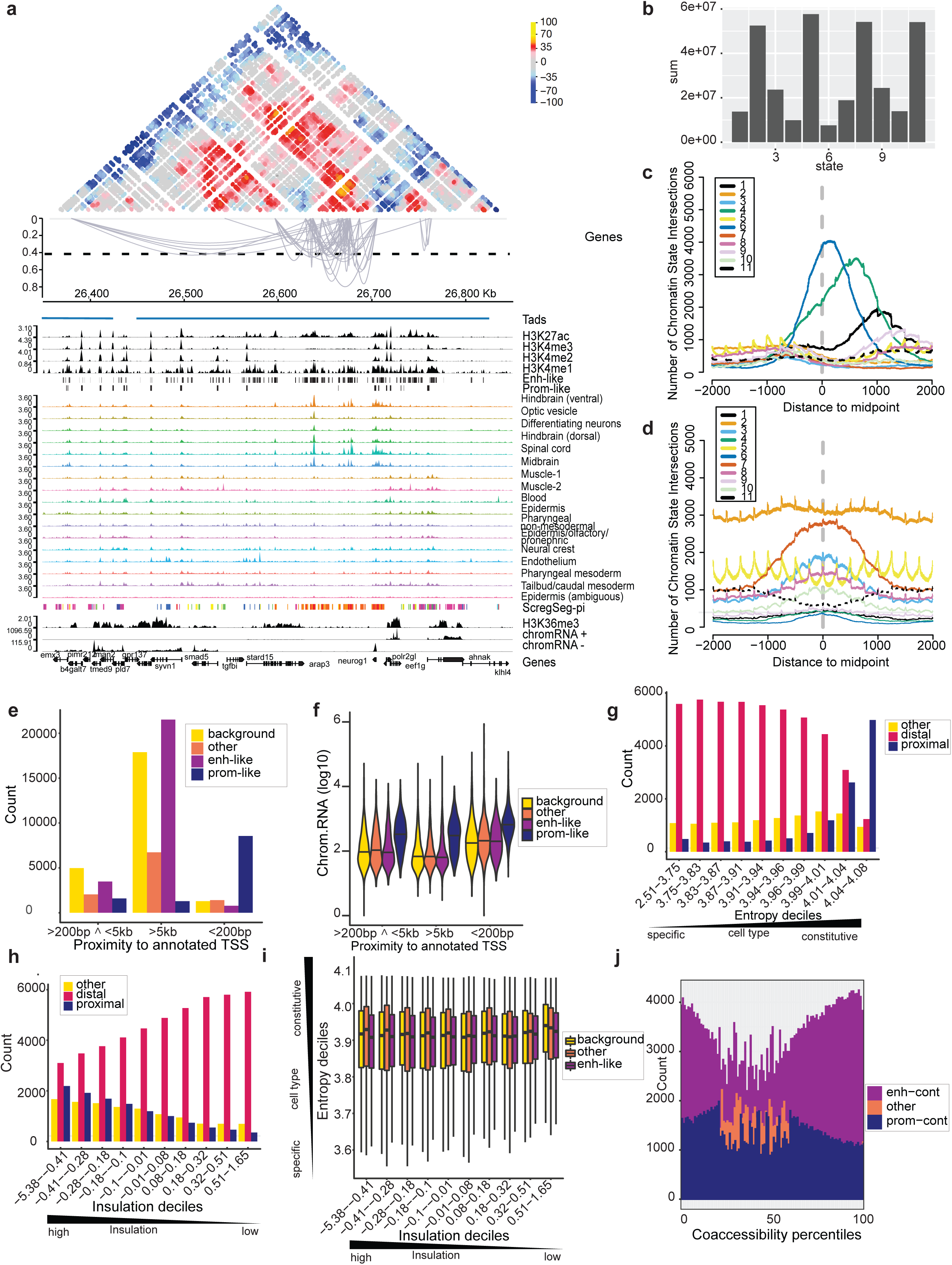
a. Browser shot as in Figure 4A except around the *neurog1* locus. b. Number of bases genome-wide covered by each histone PTM state. c. Histone PTM state coverage anchored on segment midpoints that fall +/- 200 bp from annotated TSSs. d. Histone PTM state coverage anchored on segment midpoints that fall > 200 bp from annotated TSSs. e. Number of foreground sci-ATAC-seq segments stratified by histone PTM classification types and distance to annotated TSSs. f. Log10 chromatin RNA read counts for 5 kb windows centered around intergenic foreground sci-ATAC-seq segments stratified by histone PTM classification types and distance to annotated TSSs. g. Sci-ATAC-seq foreground segment counts stratified by entropy score deciles and proximity to annotated TSS. “proximal” = <=200 bp; “other” = >200 bp & <=5 kb; “distal” = > 5 kb. h. Sci-ATAC-seq foreground segment counts stratified by insulation score deciles and proximity to annotated TSS. “proximal” = <=200 bp; “other” = >200 bp & <=5 kb; “distal” = > 5 kb. i. *In situ* Hi-C insulation scores for foreground sci-ATAC-seq segments were split into deciles and then split according to their histone PTM type. The entropy score is plotted for the three non-promoter-like histone PTM types. j. Number of segment pairs plotted for each stratification in Figure 4G.

**Supplementary figure S4.**
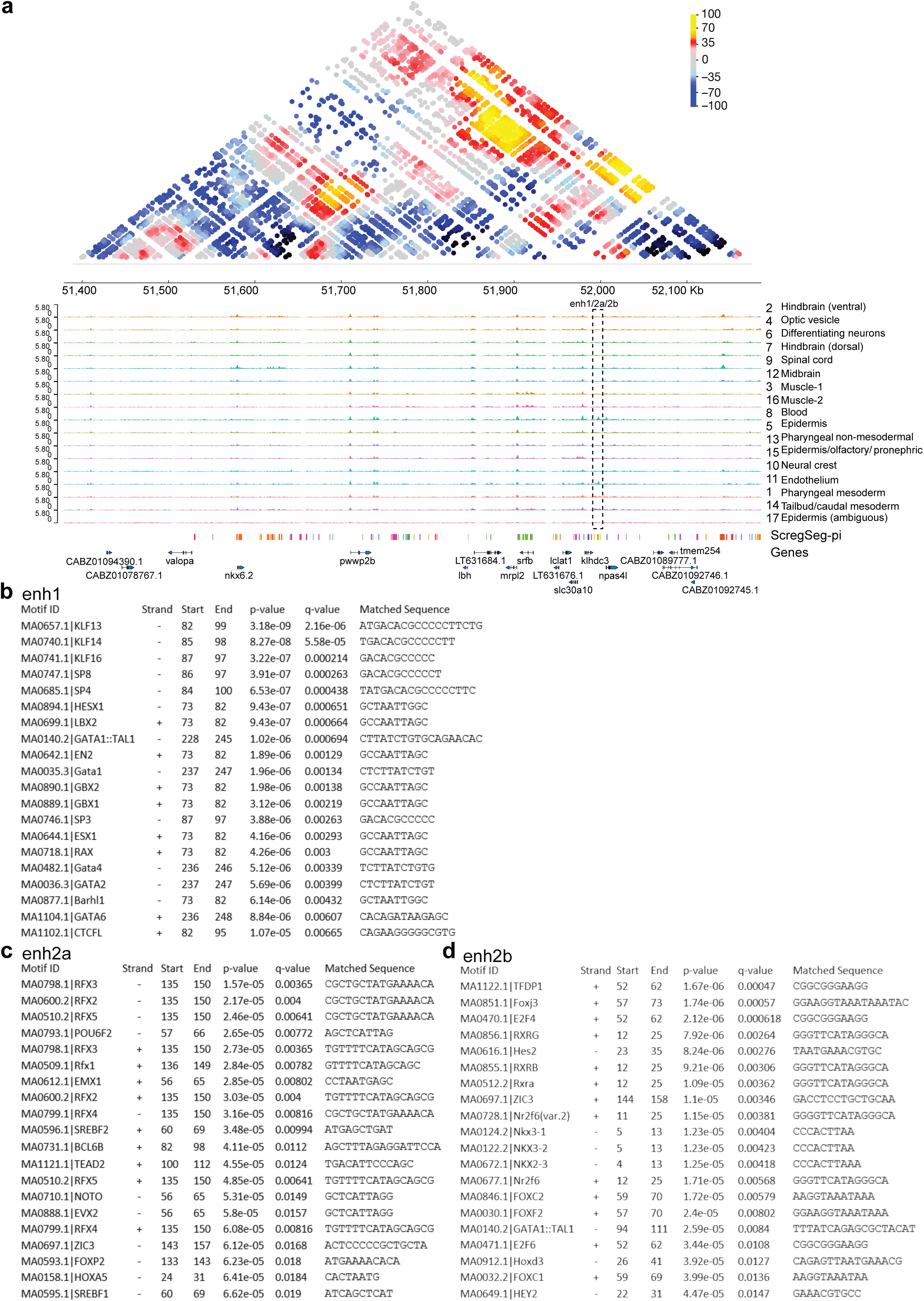
a. SHAMAN interaction scores surrounding the *npas4l* locus and it’s putative enhancers enh1, enh2a, enh2b. b. The top 20 motif matches within enh1 as detected by FIMO ^94^ using the JASPAR motif database ^56^. c. The top 20 motif matches within enh2a. d. The top 20 motif matches within enh2b.

## Supplementary tables

**Supplementary Table 1 - Enrichment scores for ZFIN-derived annotations per cluster**

**Supplementary Table 2 - Enrichment scores for scRNA-seq-derived marker genes per cluster**

**Supplementary Table 3 - Enrichment scores for ZFIN-derived annotations per ScregSeg-pi state**

**Supplementary Table 4 - Enrichment scores for scRNA-seq-derived marker genes per ScregSeg-pi state**

**Supplementary Table 5 - LogFC enrichment of genes per ScregSeg-pi state**

